# Longitudinal Developmental Trajectories Do Not Follow Cross-Sectional Age Associations in Hippocampal Subfield and Memory Development

**DOI:** 10.1101/2021.06.14.448300

**Authors:** Attila Keresztes, Laurel Raffington, Andrew R. Bender, Katharina Bögl, Christine Heim, Yee Lee Shing

## Abstract

Many cross-sectional findings suggest that volumes of specific hippocampal subfields increase in middle childhood and early adolescence. In contrast, a small number of available longitudinal studies observed decreased volumes in most subfields over this age range. Further, it remains unknown whether structural changes in development are associated with corresponding gains in children’s memory. Here we report cross-sectional age differences in children’s hippocampal subfield volumes together with longitudinal developmental trajectories and their relationships with memory performance. In two waves, 109 healthy participants aged 6 to 10 years (wave 1: *M*_Age_=7.25, wave 2: *M*_Age_=9.27) underwent high-resolution magnetic resonance imaging to assess hippocampal subfield volumes, and completed cognitive tasks assessing hippocampus dependent memory processes. We found that cross-sectional age-associations and longitudinal developmental trends in hippocampal subfield volumes were highly discrepant, both by subfields and in direction. Further, volumetric changes were largely unrelated to changes in memory, with the exception that increase in subiculum volume was associated with gains in spatial memory. Importantly, the observed longitudinal patterns of brain-cognition coupling could not be inferred from cross-sectional findings. We discuss potential sources of these discrepancies. This study underscores that children’s structural brain development and its relationship to cognition cannot be inferred from cross-sectional age comparisons.

**Highlights:**

- The subiculum undergoes volumetric increase between 6-10 years of age
- Change across two years in CA1-2 and DG-CA3 was not observed in this age window
- Change across two years did not reflect age differences spanning two years
- Cross-sectional and longitudinal slopes in stark contrast for hippocampal subfields
- Longitudinal brain-cognition coupling cannot be inferred from cross-sectional data

## 1. Introduction

A prominent branch of the quest to advance our understanding of children’s memory development attempts to identify maturational trajectories of the hippocampal network (Lee et al., 2017). This interest is fueled by the hippocampus’ key functions in both learning and remembering (Scoville and Milner, 1957) that render it a brain region of particular interest to research on individual development and school-based education. Recently developed high-resolution magnetic resonance imaging (MRI) techniques provide researchers with new tools to delineate hippocampal subfields in vivo (Bakker et al., 2008; Eldridge et al., 2005). Volumetric measures of hippocampal subfields obtained using these techniques provide the current best *in vivo* proxy for assessing maturity of intrahippocampal networks in children (Keresztes et al., 2018). In this study, we examine hippocampal subfield development and its relationship with children’s memory performance by assessing both cross-sectional age-differences and longitudinal change in hippocampal subfield volumes and memory.

### 1.1 Developmental trends inferred from cross-sectional comparisons versus longitudinal studies

In line with histological studies of monkeys (Lavenex and Lavenex, 2013), high-resolution structural MRI investigations of children have observed age-related volumetric differences in the hippocampus (for reviews, see Keresztes et al., 2018; Lee et al., 2017) suggesting that hippocampal development extends at least into early adolescence, but potentially into adulthood. Importantly, this may demonstrate a more protracted development than previous human data had indicated (Frotscher and Seress, 2007; Utsunomiya et al., 1999). This line of research also provided evidence for heterogeneous hippocampal developmental trends, previously observed using longitudinal standard resolution MRI (Gogtay et al., 2006). High-resolution MRI studies consistently reported volumetric age-differences in the dentate gyrus (DG) and cornu ammoni (CA) regions. Several studies also reported age-differences in the subiculum (SUB), but no studies found age-differences in the entorhinal cortex (EC) (reviewed in Lee et al., 2017; for more recent studies see Canada et al., 2019; Keresztes et al., 2017, 2020; Riggins et al., 2018). These results have consistently suggested volumetric increases from early middle childhood until adolescence (Daugherty et al., 2017, for an exception potentially due to specific delineation methods see 2015).

To our knowledge, three previous studies have examined longitudinal trajectories of subfield volumes in children. Critically, their findings stand in stark contrast to cross-sectional results. One early study (Tamnes et al., 2014) found volumetric decrease in all subfields measured within the hippocampus proper, except in the right CA1 from 8-21 years of age. Another study from the same lab (Tamnes et al., 2018), but with an independent sample of participants aged 8-26 years, found linear decreases in most subfields, except for CA1 with an increase until 20 followed by a decrease, as well as an increase until age 13-15 followed by decelerated decrease in the SUB. The third study (Canada et al., 2021) found volumetric increases in all subfields, but only in specific age windows and hippocampal sections (body or head) for specific subfields: DG2-4/CA3 and SUB body showing an increase between 5-6 years of age, and CA1 head showing an increase between 4-5 years of age, within the 4-8 years age-window covered in their study sample.

Sources of discrepancies across and among cross-sectional and longitudinal findings may involve differences in methods of subfield delineation (Yushkevich et al., 2015a) in scanning parameters, in age ranges, in intervals between waves in case of longitudinal studies (Canada et al., 2019), and in sample size. These potential sources of differences aside, however, theoretical accounts, simulations, and comparisons of cross-sectional and longitudinal analyses of the same data caution strongly against making inferences about change based on cross-sectional studies (Bender and Raz, 2015; Lindenberger et al., 2011; Louis et al., 1986; Nyberg et al., 2010; Pfefferbaum and Sullivan, 2015). One striking example of such discrepancy was identified by Nyberg and colleagues (2010) who showed that change across six years in frontal recruitment in healthy aging was negative despite overwhelming evidence from cross-sectional studies suggesting age-related over recruitment of frontal areas. Similarly, a recent study of hippocampal volume in a lifespan sample found no cross-sectional age-differences while longitudinal analyses of the same sample detected change (Pfefferbaum and Sullivan, 2015). Moreover, cross-sectional and longitudinal results are rarely compared within the same dataset (Canada et al., 2021, 2019; Raffington et al., 2019; Riggins et al., 2018).

### 1.2 Volume-memory associations across childhood development

An early metanalysis of 33 studies on the association between hippocampal volume and memory (van Petten, 2004) favored a “smaller is better” rather than a “bigger is better” hypothesis in childhood, adolescence, and young adulthood. However, a limited number of recent studies on associations between hippocampal subfield volumes and memory in children found positive associations between some hippocampal subfields and memory (Canada et al., 2019; Keresztes et al., 2017; Riggins et al., 2018). In addition, two of these studies revealed changes in direction of volume-memory associations across age-windows of development (Canada et al., 2019; Riggins et al., 2018). Although such shifts in the direction of volume-memory associations may be partly driven by measurement variance across the tested age-window, they do point out that volume-memory associations may be the outcome of diverse underlying mechanisms potentially changing along different trajectories across childhood. This notion additionally underlies the need for longitudinal tests of volume-memory associations.

### 1.3 The current study

Here we examine cross-sectional age-differences and longitudinal change in hippocampal subfield volumes as well as their relationship with 6-10-year-old children’s memory performance. We assessed age-differences and change across two years in aggregated hippocampal subfield volumes, in memory measures assessing function of the hippocampus as a whole or of its specific subfields, as well as in hippocampal subfield volume–memory associations across time. Based on cross-sectional age–volume associations, we hypothesized that DG-CA3, CA1-2, and potentially SUB would exhibit longitudinal volumetric increases, but that we would observe no increase in EC volume. Further hypotheses were based on prior cross-sectional findings on age-volume-memory associations (reviewed in Keresztes et al., 2018; for a newer study see Canada et al., 2019). Based on the role of DG-CA3 in pattern separation on highly similar inputs to the hippocampus (Yassa and Stark, 2011), we hypothesized that volumetric change in DG-CA3 will be specifically associated with change in the behavioral ability to discriminate highly similar memories, i.e., with mnemonic discrimination (Bakker et al., 2008; Berron et al., 2016). Findings by (Daugherty et al., 2017) additionally raised the possibility that change in DG-CA3 is potentially associated with change in associative memory. Finally, we expected that change in spatial memory, which seems to rely on overall function of the hippocampal circuitry (Burgess et al., 2002; Li and King, 2019) would be associated with change in total hippocampal volume, but not with change in specific subfield volumes.

Limited cross-sectional results from wave 1 of this study have been published (Keresztes et al., 2020; Raffington et al., 2020, 2019, 2018). In addition to our main objectives, here we also follow up on one striking effect found in these earlier investigations. Childhood stress has been associated with decreased hippocampal volume and impaired memory in adulthood (Frodl and O’Keane, 2013; Lupien et al., 2009). Animal studies have suggested that such effects, exerted through alterations in glucocorticoid secretion or sensitivity, may be strongest on DG-CA3 (Conrad, 2008; Pattwell and Bath, 2017). Based on these findings, chronically elevated levels of cortisol have been hypothesized to have negative effects on hippocampal development that may lead to memory deficits. In line with these hypothesis, Keresztes et al. (2020) found a negative cross-sectional association between hair cortisol measures and DG-CA3 volume (for a similar finding, see Merz et al., 2019). Therefore, in addition to our primary objectives, we also examined whether this cross-sectional association was also observed in wave 2 of our study, and whether changes in hair cortisol are associated with longitudinal changes in subfield volumes or memory.

## 2. Methods

### 2.1 Participants

Participants included 147 children (6.08–7.98 years, *M*_age_ = 7.19, *SD*_age_ = 0.46, 67 girls) from socioeconomically diverse families in Berlin, Germany who were enrolled in a longitudinal study (for detailed descriptions, see Keresztes et al., 2020; Raffington et al., 2018). Of them, 127 returned at wave 2, approximately two years later (8.3–10.15 years, *M*_age_ = 9.25, *SD*_age_ = 0.45, 59 girls), also described in detail elsewhere (Raffington et al., 2019). A subsample of children participated in one magnetic resonance imaging (MRI) session at both waves within three weeks of the behavioral sessions. In this study we included all children who had a high-resolution structural scan in at least one of the waves. The final sample therefore consisted of 109 children. Hippocampal subfield data available at both waves was lower (*n* = 65, 24 girls; 6.08–8.00 years, *M*_age_ = 7.26, *SD*_age_ = 0.42 at wave 1, and 8.38–10.12 years, *M*_age_ = 9.27, *SD*_age_ = 0.42 at wave 2). Completeness of data for each variable, sex and age for complete cases, as well as causes of exclusion per variable, are reported in the supplement (Table S1). Informed consent was obtained from legal guardians of all participants: Parents provided written informed consent and children provided verbal assent. The study was approved by the ‘Deutsche Gesellschaft für Psychologie’ ethics committee (YLS_012015).

### 2.2 Procedure

At wave 1, all participants were invited for two sessions, and a subsample of participants was also invited for a third session. Each session lasted approximately two hours, and included various behavioral tests, physiological measurements, and questionnaires. Session 1 and 2 took place on consecutive days. Session 3 followed within three weeks of session 2 and included a ∼35-minute-long MRI session. Of interest to the current study, participants completed an associative memory task followed by a spatial memory task at the beginning of session 1, had hair samples taken for cortisol measurement at session 2, and underwent structural MRI and completed a mnemonic similarity task at the end of session 3. At wave 2, all participants returning from wave 1 were invited to participate in two sessions, each two-hours-long. Session 1 comprised of behavioral tests and two ∼ 20-minute-long MRI sessions separated by a break, as well as sampling hair for cortisol measurement. Session 2, which followed session 1 within three weeks, comprised of only behavioral tests. Of interest, the associative memory, the spatial memory, and the mnemonic similarity tasks were all performed at session 2.

### 2.3 Volumetric measurement of hippocampal subfields

Acquisition of high-resolution MRI images of the hippocampus and surrounding mediotemporal areas, and the full process of subfields delineation on MRI images was identical across waves and is described in detail in Keresztes et al. (2020). In brief, partial field of view (FoV) proton density (PD)-weighted T2 images (FoV: 206 mm; repetition time (TR): 6,500 ms; echo time (TE): 16 ms; number of slices: 30; voxel size: 0.4 mm × 0.4 mm × 2.0 mm) acquired perpendicular to the longitudinal axis of the right hippocampus on a 3 T Siemens Magnetom TrioTim syngo MRI scanner were segmented using Automated Segmentation of Hippocampal Subfields (ASHS) (Yushkevich et al., 2015b) using a custom atlas also created using ASHS from manual segmentations with excellent reliability from earlier studies (Bender et al., 2018). This approach has been shown to be highly reliable and valid in identifying hippocampal subfield boundaries in a lifespan samples, including 6–14 year-old children (Bender et al., 2018).

We delineated three regions within the hippocampal body bilaterally – the SUB, a region including CA regions 1 and 2 (CA1-2), and a region including the dentate gyrus and CA3 (DG-CA3) – as well as the EC on 6 consecutive slices anterior to the hippocampal body (see Figure 1 in Keresztes et al., 2020). Volumetric measures for these four target regions were corrected for intracranial volume based on (Jack et al., 1989; Keresztes et al., 2017; Raz et al., 2005). Where not mentioned otherwise, the resulting adjusted volumetric measures were summed across hemispheres, and used for all analyses presented in this study. To establish reliability of manually identified boundaries of the hippocampal body, K.B. and A.K. both determined the starting and end slices, for left and right hemispheres, in a subset (n = 48) of the sample (for all, Cohen’s kappa > .82), first at wave 1. Second, before determining hippocampal body boundaries in wave 2, K.B. established reliability of boundaries determined by herself, by determining boundaries again on a subset (n = 13) of the sample from wave 1 (Cohen’s kappas > .76 for left and right starting and end slices). Then, K.B. determined hippocampal body boundaries on the whole sample of wave 2.

**Figure 1.**
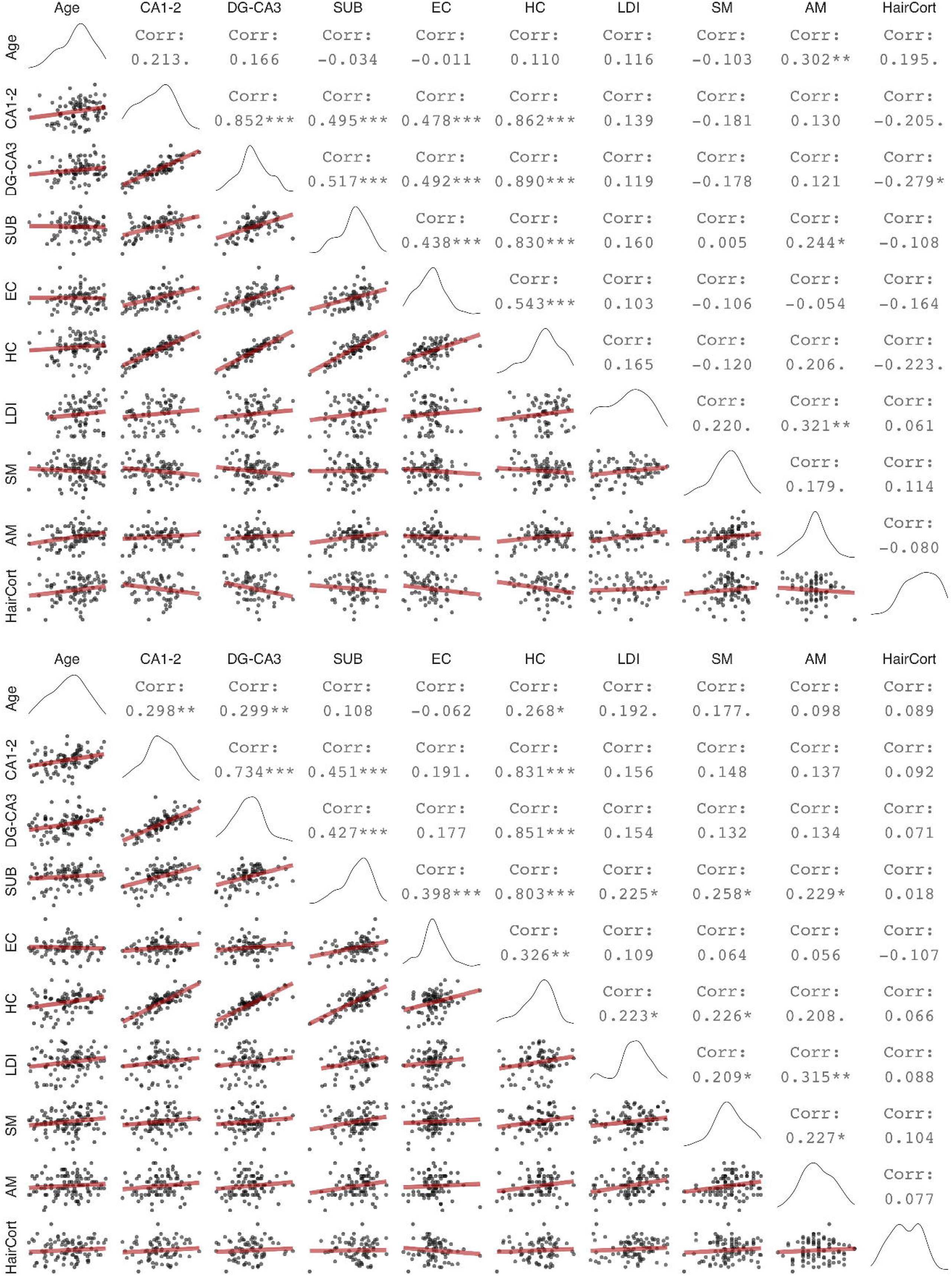
Zero order correlations between hippocampal subfield volumes and memory measures at wave 1 (upper panel) and wave 2 (lower panel). Diagonal shows variable distributions, the area under the diagonal shows scatter plots with linear regression fits (red), the area above the diagonal shows bivariate Pearson’s correlation values, with trends and significant values marked as follows: ‘.’: p < 0.1, *:p < 0.05, **:p < 0.01, ***:p < 0.001 (uncorrected for multiple comparisons). DG: Dentate gyurs, SUB: Subiculum, EC: Entorhinal cortex, HC: Hippocampus, LDI: Lure discrimination index, SM: Spatial memory, AM: Associative memory, HairCort: Hair cortisol

### 2.4 Memory measures

We administered three memory tasks to assess hippocampal contributions to memory: a mnemonic similarity task, a spatial memory task, and an associative memory task. All three were computerized tasks performed on desktop computers with 21.5-inch screens.

The mnemonic similarity task was based on (Kirwan and Stark, 2007), a continuous recognition memory task assessing participants’ ability to discriminate memories of highly similar stimuli. Participants saw 162 pictures depicting everyday objects, presented sequentially. Critically, 48 pictures were repeated after a delay of 2–14 intervening trials. Twenty-four of the pictures were repeated exactly, whereas 24 pictures were repeated with a slightly different lure picture of an identical object. For each trial the children’s task was to identify pictures as “old” (exact repetition), “similar” (lure repetitions), or “new” (new items). We calculated a lure discrimination index as the proportion of “similar” responses to lure repetitions minus the proportion of “similar” responses to new items reflecting participants’ ability to discriminate between highly similar memories. More details on this task are given in (Keresztes et al., 2020).

The spatial memory task was based on methods originally reported by Kessels et al. (2007), and assessed memory for items and item-location associations. In brief, participants encoded locations of 15 sequentially presented line drawings of everyday objects on a 6 × 6 grid. After a short delay, they performed a recognition task using the same pictures of studied objects randomly intermixed with 15 pictures of new objects. For each correctly recognized object, we asked them to point to the location of the given picture in the grid during encoding. As a measure of spatial memory, we used the percentage of correctly indicated locations for the 15 old items. For a detailed description of the task see (Raffington et al., 2018).

Participants also completed a paired-associate memory task, with an incidental encoding phase followed by two retrieval phases, one at a short (< 2 mins) delay and one after approximately 24h to assess associative memory performance across different time spans. At encoding, 34 pairs of German nouns (for the list of words see Table S2 in the Appendix) were presented to participants sequentially. Each word pair was presented simultaneously both in visual (on the computer screen) and auditory modalities (via loudspeaker). In each trial, following a fixation cross (500 ms), a cue word appeared in the middle of the screen for 2 secs, then the target word appeared on the screen below the cue word; both cue and target words remained on screen for an additional 4 secs. Participants then saw a question mark on the screen for 15 secs, and their task was to decide whether the two words are related to each other or not. The two retrieval phases were identical, both came as a surprise to participants, and both included different halves of the word pairs encoded. The retrieval phase consisted of a cued recall block followed by a stem-cued recall block. Trials in cued recall consisted of a cue presented both visually and auditorily, with the visually presented cue remaining on screen for 15 secs or until the participant pressed a response button on a keyboard. Then the experimenter asked the participant to report the answer verbally. The experimenter typed in the answer via the same keyboard and then advanced the experiment to the next trial. Trials in the stem cued recall block were identical to trials in the cued recall block but the cue was appended with the first two phoneme of the target (e.g. ‘Gi’ for Giraffe, and ‘Kr’ for “Kreide”/chalk). Because performance on cued recall was close to floor on the second retrieval test performed a day after encoding, and because stem-cued recall is known to be less dependent on hippocampal processing, in this study we only used cued recall performance on the first retrieval test. As a measure of associative memory, we used percentage of targets correctly recalled.

### 2.5 Hair cortisol measurement

Details of methods for hair cortisol assessment are given in Keresztes et al. (2020). In brief, the Dresden LabService GmbH, Germany, analyzed the first 3 cm long segment of a ∼1 mm thick strand of hair taken as close to the skull as possible (Gao et al., 2013). Cortisol concentrations (pg/mg), log-transformed for parametric statistics, were used as a cumulative estimate of hormones secreted over 3 months prior to sampling (Stalder et al., 2016). Wave 1 and wave 2 hair cortisol samples were assayed as separate batches. Thus, mean change in hair cortisol levels between wave 1 and 2 may be confounded with batch effects. Nevertheless, we could examine variance in hair cortisol change and change-change associations, assuming that any batch effect affects samples to the same degree.

### 2.6 Cross-sectional and longitudinal assessment

To assess cross-sectional age-differences, we linearly regressed variables of interest on age. We assessed bivariate associations between hippocampal subfield volumes, memory measures and hair cortisol cross-sectionally using Pearson’s zero-order correlations. To assess longitudinal change across two waves, we adopted latent change score (LCS) structural equation models (Kievit et al., 2018; McArdle and Nesselroade, 1994), using the lavaan package (Rosseel, 2012) in R (R Core Team, 2019). The LCS models produced three parameter estimates of particular interest: (1) mean change from wave 1 to wave 2, (2) variance in change, and (3) the covariance between the intercept (wave 1 values) and change. After fitting univariate LCS models to assess change in each of the variables (see Figure 2a), we employed a bivariate LCS approach (Figure 4a) to assess pairwise change–change associations between variables. Age at wave 1 was included as a covariate in all models, covarying with change, and predicting wave 1 intercept. For estimation of LCS model parameters, hippocampal subfield volume values were divided by 100 to match the scale of other variables in the analyses.

**Figure 2.**
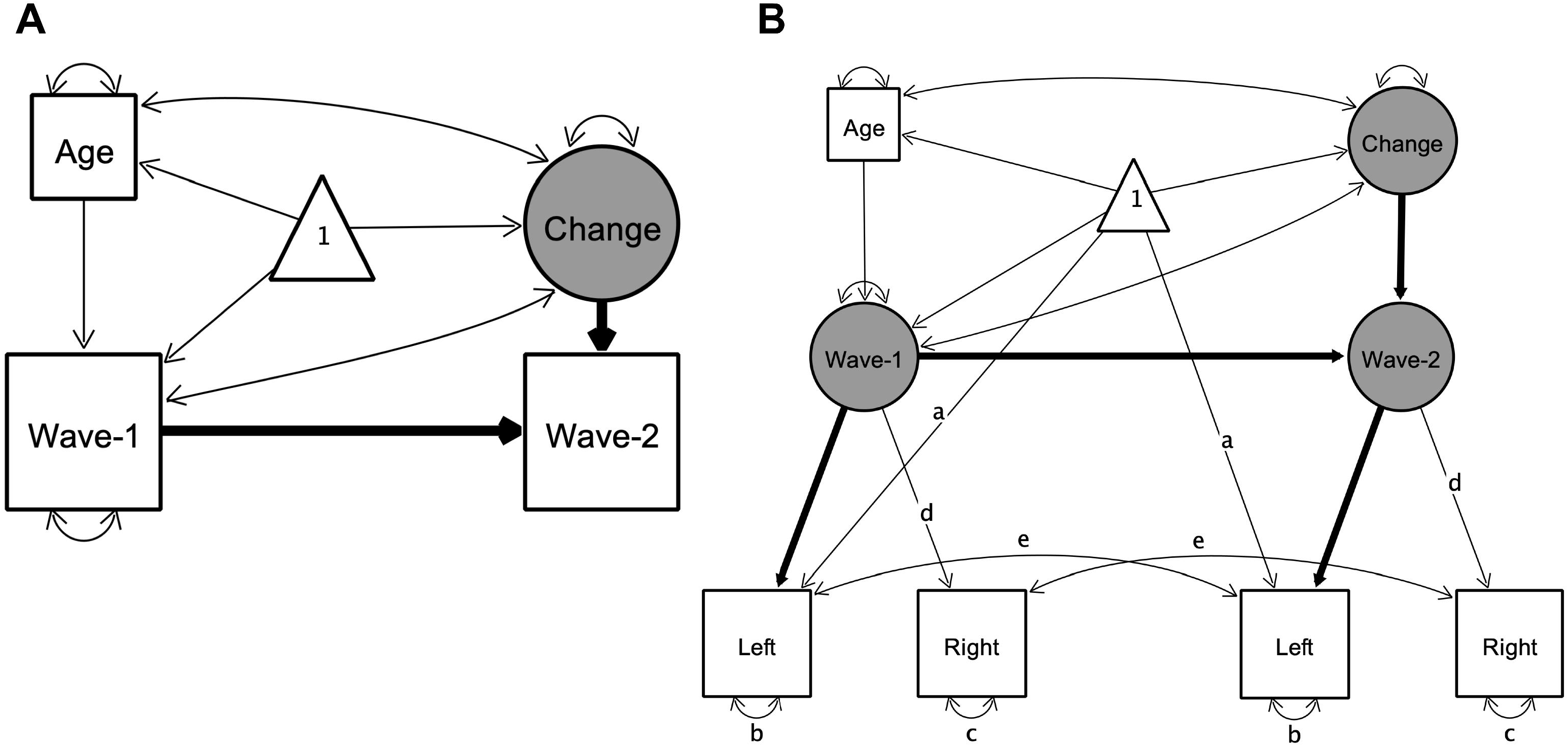
Illustration of univariate latent change score models used to investigate change in variables of interest. Rectangles, circles, and triangles represent indicator variables, latent variables, and means, respectively. Estimated variances, covariances and regression paths are shown as thinner solid lines. Thicker solid lines represent path values fixed at 1. Age represents age at wave 1. In (A) used to model change in all variables of interest, wave-1 and wave-2 represent observed values of a given variable at each wave. In bilateral indicator univariate LCS models (B), used as an additional model of change in hippocampal subfield volumes, latent wave 1 and wave 2 estimates are modeled by observed left and right hemispheric volumetric measures. Identical path names denoted by (a–e) represent estimated parameters that are constrained to be equal to each other to ensure measurement invariance (Raz et al., 2005). Difference in χ^2^ fit statistics indicated that lifting these constraints did not improve model fit, except for EC. For EC letting indicator variances of left (b) and right (c) differ across waves improved model fit, therefore we lifted these constraints in that model (see Bender and Raz, 2015). Parameter estimates of interest are shown separately in Table 1 for model (A) and Table S3 for model (B) for better readability of the figure. Figures created by Onyx, version 1.0–1010 (von Oertzen et al., 2015).

Volumetric measures acquired using MRI are not free of measurement error (Karch et al., 2019; Maclaren et al., 2014; Madan and Kensinger, 2017), therefore in addition to LCS models with single bilateral volumetric indicators, we also tested univariate LCS models of change in hippocampal subfields where left *and* right hemispheric volumes served as dual observed indicators for the volumetric latent variables at each wave (Figure 2b and 4b). These models thus separate error variance from construct variance and establish measurement invariance over time. We used these models – henceforth referred to as bilateral indicator univariate LCS models (Kievit et al., 2018) of the hippocampal subfield volumes – to replicate findings of the univariate LCS models.

All models were computed using maximum likelihood estimation implemented in lavaan. To evaluate model fit, we used standard goodness-of-fit indices: the root mean square error of approximation (RMSEA) and the comparative fit index (CFI). Models were considered a good fit with a RMSEA < 0.08 and CFI > 0.95 (Kline, 2015). The difference in χ^2^ fit statistics was used to compare nested models, with the degrees of freedom being the difference in the number of free parameters. The threshold for statistical significance was *alpha =* 0.05. In addition, we calculated 95% confidence intervals for parameter estimates using 1000 bootstrapped resamples.

## 3. Results

### 3.1 Age-differences in hippocampal subfield volumes were stable across time points but did not match longitudinal change in hippocampal subfield volumes over time

The pattern of cross-sectional associations in wave 2 was similar to that observed in wave 1 (Keresztes et al., 2020, see also Figure 1). CA1-2, DG-CA3, and total HC volume correlated positively, and significantly with age, whereas neither SUB nor EC showed significant associations with age. However, these patterns were at odds with change observed within individuals. Latent change score models (see Figure 2a for a graphical depiction of univariate LCS model specifications; and Table 1 for model fit and parameter estimates) indicated significant, positive mean volumetric changes for SUB and EC, as well as non-significant negative mean volumetric change in CA1-2, and non-significant positive mean change in DG-CA3 and in HC. Confidence intervals calculated from bootstrapped samples (see Table S4), provided support for the robustness of the significant associations. Importantly, bilateral indicator univariate LCS models of the hippocampal subfield volumes (see Figure 2b and Table S3) also replicated these results. Discrepancies between cross-sectional and longitudinal slope estimates are visualized in Figure 3.

**Table 1.** Key parameter estimates in univariate latent change score models for all variables of interest.

|  | Parameter estimates |  |  |  |  |  |  |  |
| --- | --- | --- | --- | --- | --- | --- | --- | --- |
|  | M<sub>change</sub> |  | Var<sub>change</sub> |  | β<sub>Age-at-wave 1→wave 1</sub> |  | Cov<sub>change-wave 1</sub> |  |
|  | PE (SE) | Δχ<sup>2</sup>(1) | PE (SE) | Wald statistic<sup>a</sup> | PE (SE) | Δχ<sup>2</sup>(1) | PE (SE) | Δχ<sup>2</sup>(1) |
| *Hippocampal subfields* |  |  |  |  |  |  |  |  |
| CA1-2 | -0.08 (0.05) | 2.45 | 0.172 (0.031) | 5.6*** | 0.307 (0.153) | 3.94* | -0.094 (0.035) | 9.20** |
| DG-CA3 | 0.015 (0.065) | 0.06 | 0.288 (0.051) | 5.59*** | 0.294 (0.222) | 1.74 | -0.204 (0.066) | 12.70*** |
| SUB | 0.25 (0.088) | 7.66** | 0.533 (0.094) | 5.67*** | -0.01 (0.277) | 0.001 | -0.417 (0.109) | 22.22*** |
| EC | 0.143 (0.047) | 8.55** | 0.154 (0.027) | 5.73*** | -0.007 (0.14) | 0.003 | -0.033 (0.027) | 1.55 |
| Total HC | 0.192 (0.189) | 1.02 | 2.5 (0.451) | 5.55*** | 0.577 (0.572) | 1.01 | -1.892 (0.498) | 21.63*** |
| *Memory measures* |  |  |  |  |  |  |  |  |
| LDI | 0.101 (0.019) | 24.11*** | 0.023 (0.004) | 5.78*** | 0.044 (0.050) | 0.76 | -0.018 (0.003) | 73.23*** |
| SM | 0.084 (0.017) | 23.02*** | 0.029 (0.004) | 7.23*** | -0.036 (0.033) | 1.17 | -0.013 (0.003) | 8.375** |
| Cued recall | 0.139 (0.019) | 43.36*** | 0.036 (0.005) | 7.03*** | 0.116 (0.037) | 9.35** | -0.013 (0.003) | 22.67*** |
| *Hair cortisol*<sup>b</sup> |  |  | 0.761 (0.119) | 6.39*** | 0.337 (0.239) | 1.95 | -0.627 (0.117) | 64.67*** |

**Note.** M: mean, Var: variance, PE (SE): parameter estimate (standard error), DG: dentate gyrus, SUB: subiculum, EC: entorhinal cortex, HC: hippocampus. SM: Spatial memory, LDI: Lure discrimination index. Model fit was perfect for all models, $\chi^2=0$ , RMSEA=0, CFI=1. Estimates for error variances, as well as for indicator variable means are not presented. \*: $p < 0.1$ , \*\*: $p < 0.05$ , \*\*\*: $p < 0.01$ , \*\*\*\*: $p < 0.001$ , uncorrected for multiple comparisons. Confidence intervals calculated from bootstrapped samples provided support for the robustness of all associations significant at $p < .05$ . <sup>a</sup>: For Var<sub>change</sub> estimates, the difference in $\chi^2$ could not be obtained because the models did not converge without this parameter. Therefore, the Wald statistic was used instead. <sup>b</sup>: Due to potential batch effects, mean change was not interpreted in this model. Units are mm<sup>3</sup>/100 for hippocampal subfields, % for memory measures, and log(pg/mg) for hair cortisol.

**Figure 3.**
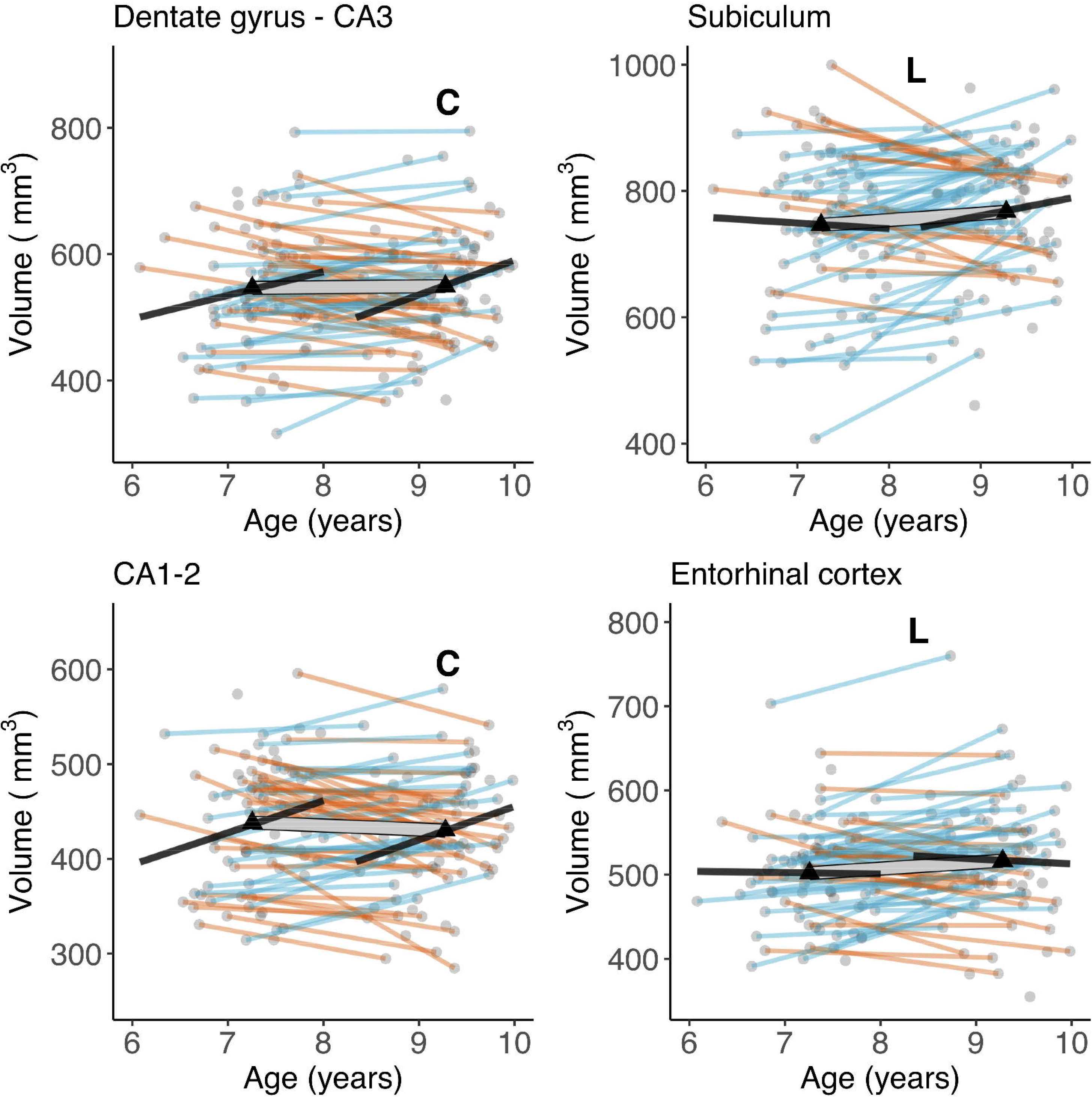
Cross-sectional age-differences and longitudinal age-trends in hippocampal subfield volumes. Dots represent individuals, triangles represent mean volume at each wave. Black lines show regressions of volume on age cross-sectionally at each wave. The thick transparent white line represents mean change, thinner colored lines represent individual change with orange and blue lines showing a decrease or increase in volume, respectively. Bold letters ‘C’ and ‘L’ represent significant cross-sectional age-difference at a given wave, and significant mean change across waves, respectively.

We observed significant variance in volumetric change in all hippocampal subfields (see Table 1 and Figure 3). As shown in Figure 3, the number of individuals showing volumetric increase (versus decrease) differed across subfields, with 27 (42%), 34 (52%), 46 (71%), and 42 (65%) of 65 participants showing volumetric increase in CA1-2, DG-CA3, SUB, and EC, respectively. Volumes at wave 1 were also significantly negatively associated with volumetric change for all subfields, except for EC. Removing age as a covariate from LCS models did not change the observed pattern of results.

### 3.2 Cross-sectional age-differences in memory measures were stable across time points and agreed with change in memory measures over time

Except for spatial memory at wave 1, memory measures and age were positively associated at both waves, although most of these associations did not significantly differ from zero (Figure 4). LCS models on memory measures (see Table 1) showed a significant linear mean change over time for all three memory measures. We observed significant variance in change across the three tasks, with 61%, 74%, and 71% of participants showing an increase in spatial memory, lure discrimination and associative memory respectively. Wave 1 performance on all three tasks was negatively associated with change in performance across the two waves. Again, removing age as a covariate from LCS models did not change the observed pattern of results.

**Figure 4.**
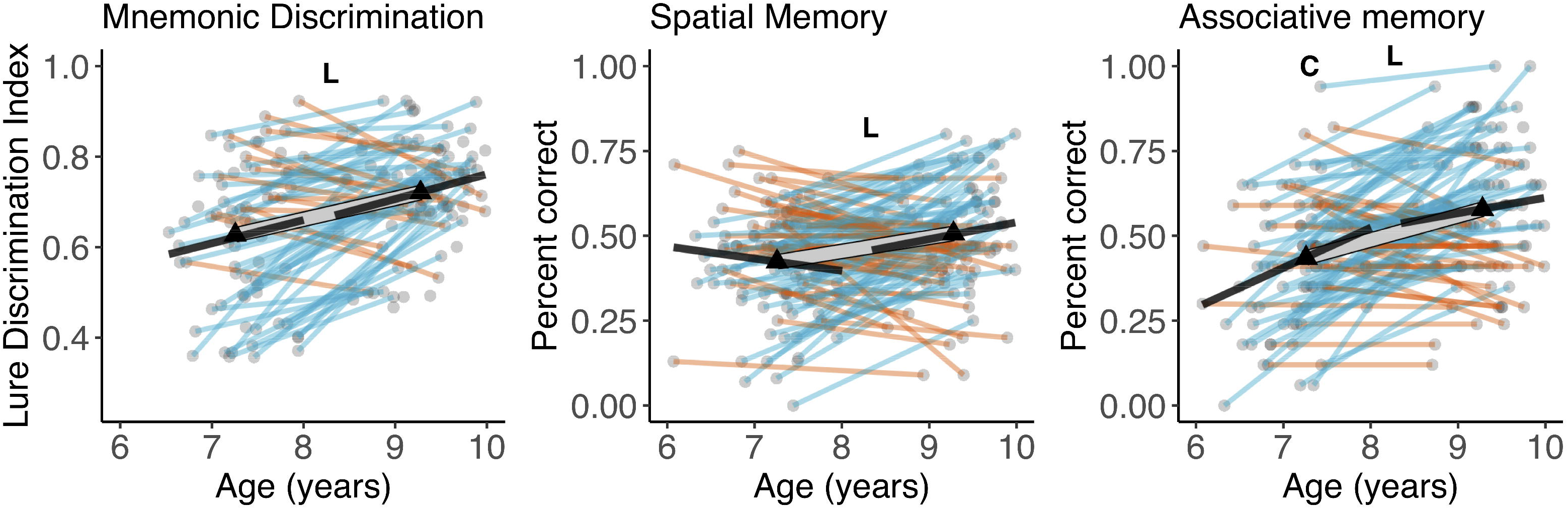
Cross-sectional age-differences and longitudinal age-trends in memory measures. Dots represent individuals, triangles represent mean volume at each wave. Black lines show regressions of volume on age cross-sectionally at each wave. The thick transparent white line represents mean change, thinner colored lines represent individual change with red and green lines showing a decrease or increase in volume, respectively. Bold letters ‘C’ and ‘L’ represent significant cross-sectional age-difference at a given wave, and significant mean change across waves, respectively.

### 3.3 No cross-sectional age-differences in hair cortisol and significant variance in change across participants

In both wave 1 and wave 2, we observed a non-significant positive association between age and hair cortisol (Figure 1). Univariate LCS models indicated significant variance in change. and a significant negative association between wave 1 hair cortisol concentrations and change in hair cortisol concentrations over time (Table 1). Note that because of the batch effects in hair cortisol measures (described in section 2.5) we cannot infer whether the latter association reflects that higher baseline hair cortisol levels are associated with a negative or a less positive change.

### 3.4 Consistency of cross-sectional associations between hippocampal subfields, memory measures, and hair cortisol at wave 1 and wave 2

Zero-order correlations between volumetric measures of hippocampal subfields and memory measures (Figure 1) showed that SUB was the only hippocampal subfield significantly associated with memory at either wave. In wave 1, SUB correlated positively and significantly with associative memory, and in wave 2 it significantly correlated with all three memory measures. Thus, only one association – between associative memory and SUB volume – was consistent in both waves. In the model of total HC, we observed significant positive associations between HC and lure discrimination and spatial memory at wave 2, as well as a positive trend at both waves between HC and associative memory. The significant negative correlation found at wave 1 between hair cortisol and DG-CA3 volume was absent at wave 2, and no other correlations with hair cortisol were significant in either waves.

### 3.5 Longitudinal associations between hippocampal subfields, memory measures, and hair cortisol

Next, we tested for potential longitudinal associations between performance on memory measures and volumetric measures in hippocampal subfields using bivariate LCS models. Altogether, we ran 15 LCS models, 4 subfield volumetric measures plus volume of total hippocampal body *×* 3 memory measures, and extracted parameter estimates for change–change covariances as well as covariances between memory measures at wave 1 and change in hippocampal subfield volumes, and between hippocampal subfield volumes at wave 1 and change in memory (see Figure 5a for bivariate LCS model specification, and Table 2 for model fit and parameter estimates).

**Figure 5.**
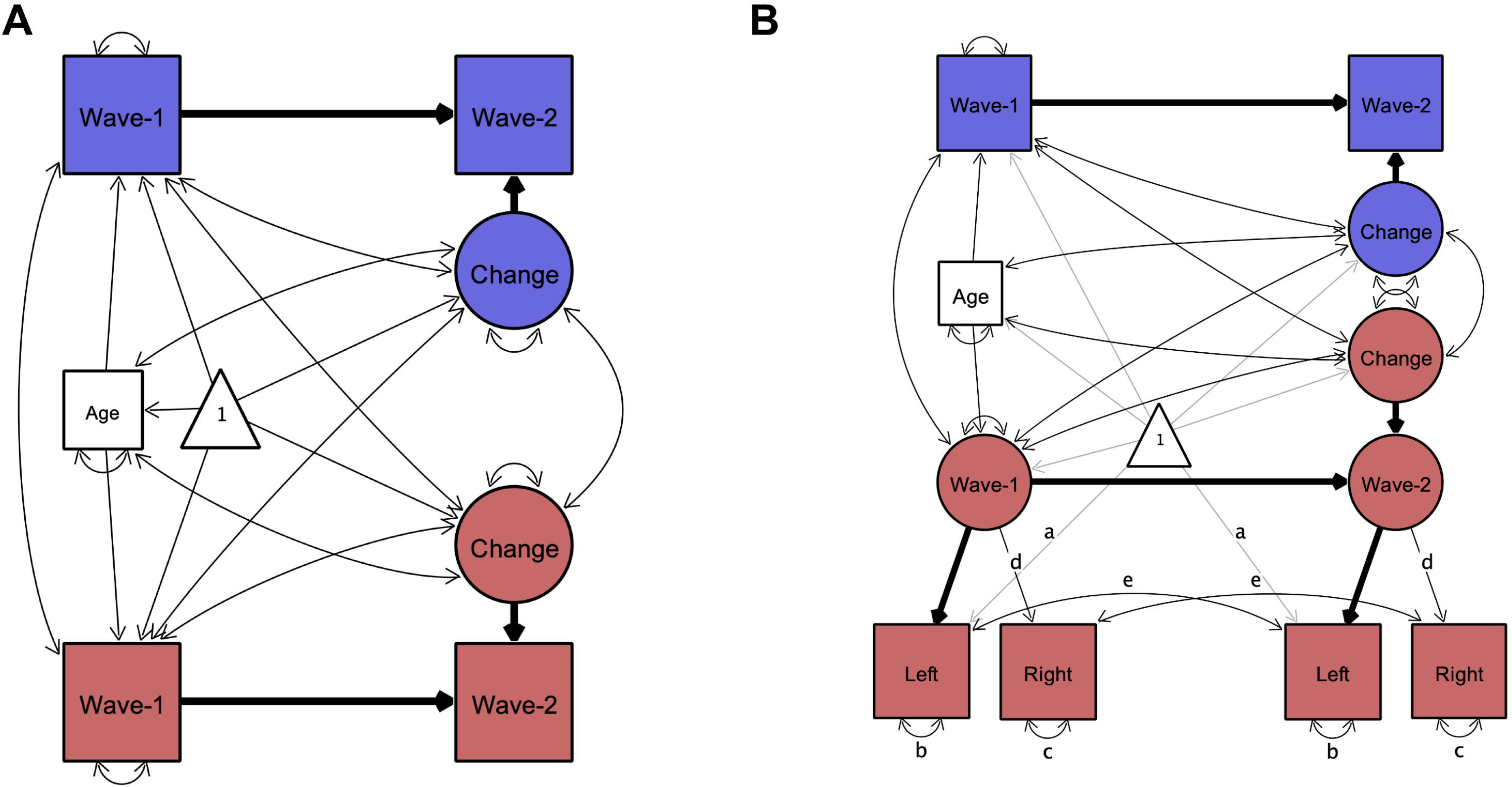
Illustration of bivariate latent change score models used to assess change-change association between hippocampal subfield volumes and memory measures. Rectangles, circles, and triangles represent indicator variables, latent variables, and means, respectively. Estimated variances, covariances and regression paths are shown as thinner solid lines. Thicker solid lines represent path values fixed at 1. Age represents age at wave 1. Blue and red colors used to distinguish between memory and hippocampal subfield variables respectively. In (A) wave-1 and wave-2 represent observed values of both variables at each wave. In (B), latent wave 1 and wave 2 estimates of hippocampal subfields are modeled by observed left and right hemispheric volumetric measures. Identical path names denoted by (a–e) represent estimated parameters that are constrained to be equal to each other to ensure measurement invariance (Raz et al., 2005). Paths representing means are shaded for better visibility. Parameter estimates of the change-change covariance between hippocampal subfields and memory measures are shown separately in Table 2 for model (A) and in Table S5 for model (B) for better readability of the figure. Figures created by Onyx, version 1.0–1010 (von Oertzen et al., 2015).

**Table 2.** Model fit, and parameter estimates for covariance between change in hippocampal subfields and change in memory measures in bivariate latent change score models.

|  | **LDI** |  | **Spatial memory** |  | **Cued recall** |  |
| --- | --- | --- | --- | --- | --- | --- |
|  | **PE (SE)** | **$\Delta\chi^2(1)$** | **PE (SE)** | **$\Delta\chi^2(1)$** | **PE (SE)** | **$\Delta\chi^2(1)$** |
| **CoV<sub>change-change</sub>** |  |  |  |  |  |  |
| **CA1-2** | 0 (0.008) | 0.002 | 0.005 (0.008) | 0.395 | 0.005 (0.011) | 0.225 |
| **DG-CA3** | -0.004 (0.012) | 0.119 | 0.012 (0.011) | 1.225 | -0.003 (0.014) | 0.043 |
| **SUB** | 0.002 (0.015) | 0.027 | 0.03 (0.015) | 4.395* | 0.006 (0.02) | 0.096 |
| **EC** | -0.001 (0.009) | 0.005 | -0.002 (0.008) | 0.057 | -0.006 (0.01) | 0.313 |
| **HC** | -0.004 (0.033) | 0.012 | 0.047 (0.032) | 2.199 | 0.01 (0.041) | 0.055 |
| **CoV<sub>volume at wave 1 – change in memory</sub>** |  |  |  |  |  |  |
| **CA1-2** | -0.005 (0.011) | 0.181 | 0.006 (0.011) | 0.301 | 0.001 (0.013) | 0.008 |
| **DG-CA3** | -0.009 (0.017) | 0.283 | 0.002 (0.015) | 0.026 | -0.005 (0.019) | 0.083 |
| **SUB** | -0.016 (0.020) | 0.668 | -0.008 (0.019) | 0.197 | -0.005 (0.023) | 0.038 |
| **EC** | -0.005 (0.011) | 0.193 | 0.001 (0.01) | 0.004 | 0.017 (0.012) | 2.181 |
| **HC** | -0.032 (0.042) | 0.588 | 0.001 (0.039) | 0.000 | -0.011 (0.048) | 0.057 |
| **CoV<sub>memory at wave 1 – change in volume</sub>** |  |  |  |  |  |  |
| **CA1-2** | 0.005 (0.009) | 0.283 | 0.01 (0.008) | 1.602 | -0.003 (0.009) | 0.085 |
| **DG-CA3** | 0.012 (0.012) | 1.028 | 0.01 (0.01) | 0.994 | 0.009 (0.012) | 0.531 |
| **SUB** | 0.011 (0.015) | 0.485 | 0.011 (0.014) | 0.629 | 0.001 (0.017) | 0.006 |
| **EC** | 0.002 (0.01) | 0.034 | 0.016 (0.007) | 4.788* | 0.001 (0.008) | 0.004 |
| **HC** | 0.031 (0.034) | 0.846 | 0.034 (0.03) | 1.353 | 0.006 (0.035) | 0.034 |

**Note.** M: mean, Var: variance, PE (SE): parameter estimate (standard error), DG: dentate gyrus, SUB: subiculum, EC: entorhinal cortex, HC: hippocampus. LDI: Lure discrimination index. Model fit was perfect for all models, $\chi^2=0$ , RMSEA=0, CFI=1. Estimates for error variances, as well as for indicator variable means are not presented. \*: $p < 0.05$ , uncorrected for multiple comparisons.

The only significant change-change association was a positive one between change in SUB volume and change in spatial memory performance (*p* = .036). However, a confidence interval calculated from bootstrapped samples [−0.001,0.065] did not provide support for the robustness of this association. In addition, we observed a significant positive association between performance on the spatial memory task at wave 1 and change in EC (*p* = .029). A confidence interval from bootstrapped samples [0.001,0.03] provided additional support for the robustness of this association. No other longitudinal parameter estimates of interest were significant. Additional bivariate LCS models detected no association between change in hair cortisol and change in hippocampal subfield volumes. Importantly, bivariate LCS models including bilateral indicator univariate LCS models for the hippocampal subfield volumes (see Figure 5b), replicated the observed pattern of results (see Table S5), that was also unaltered when age was removed as a covariate from the models.

### 3.6 Sex effects

We tested for sex differences as well as for age *×* sex interactions in all variables of interest, using independent samples t-tests and multiple linear regressions, respectively. We found a significant difference in EC volume – with girls having larger volumes – in both wave 1 (*t*(82) = 2.1, *p* = .04), and wave 2 (*t*(83) = 4.06, *p* < .001). No other sex effects were significant, nor any sex *×* age interactions. Including Sex in LCS models of EC did not change the pattern of results reported in 3.1 and 3.4.

### 3.7 Additional control and power analyses

We performed additional independent sample t-tests to rule out the possibility that selective attrition between waves may underlie, at least in part, the discrepancies between estimates of cross-sectional age-differences and longitudinal change. For instance, if dropouts have lower or higher values compared to ongoing participants, we could observe longitudinal changes even if there is no actual change or miss out on detecting longitudinal change. Because we also had participants who participated in both waves but were scanned only at wave 2, we defined dropouts for analyses both forward (wave 1 data present and wave 2 data missing) and backward (wave 2 data present and wave 1 data missing) in time. For all volume and memory variables, for both forward and backward dropouts, we performed Welch t-tests comparing dropouts with non-dropouts. Neither of these analyses yielded significant effects (all *t*s < .97, all *p*s > .34), suggesting that potential differences between dropouts and non-dropouts were not affecting longitudinal estimates of change in any variables of interest.

In addition, we also tested whether dropouts differed from non-dropouts on sex or age. Forward dropouts were significantly older than non-dropouts, *t*(8.7) = 4.00, *p* = .003. We found no other age or sex differences between dropouts and non-dropouts (See section S1.1 of the Supplement for the same analyses performed separately for each variable of interest separately).

To rule out a limit on power to detect otherwise existing associations in our relatively small sample, we performed post-hoc power analyses for univariate LCS models using the RAMpath R package (Zhang et al., 2015; Zhang and Liu, 2018), based on the original RAMpath software (Boker et al., 2002), relying on lavaan (Rosseel, 2012). This analysis indicated that LCSs in our longitudinal sample, given an *α* = 0.05, had a power of .99 to detect a mean volumetric change in DG-CA3 and CA1-2; this was equivalent to the smallest cross-sectional slope for the same regions across the two waves (slope of CA1-2 on Age at wave 1 = 0.34). Because longitudinal slopes may indeed be smaller than cross-sectional slopes, we drew power curves plotting power against sample size as a function of longitudinal slope estimates. This analysis (see Figure S1) indicated that our LCS model’s power decreased below the conventionally accepted .8 value when the estimated slope was below .19, meaning that we may have missed existing effects of change if these were below an annual change of 19 mm^3^.

## 4. Discussion

The two most important highlights from our study are (1) we found longitudinal changes that had not been predicted based on cross-sectional data, and (2) most of our hypotheses based on cross-sectional age associations were not supported by longitudinal data. Crucially, for hippocampal subfields, we found a cross-sectional trend for age-related differences in CA1-2 at wave 1 and significant age-related differences in CA1-2 and DG-CA3 at wave 2, coupled with no differences in SUB and EC. This pattern of results is partly in line with the available evidence for cross-sectional age-differences in childhood (Canada et al., 2019; Keresztes et al., 2018; for a review, see: Lee et al., 2017; Riggins et al., 2018). Together, these findings would suggest an ongoing development of hippocampal subfields potentially lasting into adolescence.

In sharp contrast, our hypothesis that DG-CA3 and CA1-2 volumes increase over time was not supported by our longitudinal analysis. Rather, we observed significant positive changes for SUB and EC. We observed substantial individual variation in change over time across all variables except for EC. Some of these variations were due to a negative association between initial values and change. Despite high variability across individuals, we did not find strong change-change associations between subfield volume and memory. Importantly, we found no support for the hypothesis that change in DG-CA3 volume is associated with change in the ability to discriminate highly similar memories. Instead, our data showed an association between change in SUB volume and change in performance on a spatial memory task, although this weak effect should be dealt with caution. That said, it is worth noting that this effect was not predicted based on the structure–memory associations observed at wave 1, but is somewhat consistent with associations between SUB volume and memory found at wave 2. In addition, we found a positive association between spatial memory performance at wave 1 and change in EC volume. Because this was also an unpredicted and weak association we refrain from interpreting this specific association until further replication. Finally, at wave 2 we did not replicate the striking negative association between DG-CA3 volume and hair cortisol observed at wave 1 (Keresztes et al., 2020). Together with an earlier finding of negative association between DG-CA3 volume and hair cortisol (Merz et al., 2019) in a sample of children aged 5-9 years (M = 7.03) this result may speak to a sensitive period for cortisol effects on hippocampal subfields. However, given the limited sample sizes of both studies, this speculation needs to withstand further scrutiny.

### 4.1 Potential factors driving the disagreement between cross-sectional age-associations and longitudinal trends

Below, we first discuss how these critical discrepancies between cross-sectional and longitudinal data may emerge, and their implication for research on neural and cognitive development. Then, we consider the implications of our specific findings for research on the neurocognitive development of memory, and in particular hippocampal subfield development.

Differences between cross-sectional and longitudinal results are commonly attributed to a multitude of factors, as they stem from different studies with varying methodologies. The strength in our approach here is that we used identical methods across waves, hence effects of discrepant methodological details including delineation of subfields, the choice of volumetric measure, and delays between waves (Canada et al., 2020) were not relevant for the current study. Moreover, the cross-sectional age span and the interval between the two waves of the study were both approximately 2 years, allowing us to assess longitudinal and cross-sectional slopes on equal timescales, which rarely is the case in a longitudinal study. Second, given the small age-range of the samples collected at each wave, it is unlikely that cohort effects (Raz and Lindenberger, 2011) drive the observed age-differences in DG-CA3 and CA1-2. Third, there were no selective dropouts that could have led to disagreement between cross-sectional age-differences and longitudinal estimates of change (Nyberg et al., 2010). Although we did not find differences in memory or volumetric measures between dropouts and non-dropouts, we did find that dropouts after wave 1 were older than ongoing participants, and for some variables (see Supplement S1.1) predominantly boys. However, these differences can hardly explain the effects found: We only found sex differences in volume of EC, with boys having smaller volumes than girls. Given that wave 1 volumes were negatively associated with change for all subfields, if sex differences in dropouts had an effect, if any, on observed change in EC, this should have been an attenuation of the observed increase. In addition, because age was included as a covariate in our LCS models estimating change, age differences between dropouts and non-dropouts cannot drive the observed effect of change. That said, based on these post-hoc tests, we cannot fully rule out that we missed out on any effects, had we no dropouts at all. Fourth, using post-hoc power calculations we have also shown that despite the relatively low sample size, our study was well powered to detect the magnitude of change in DG-CA3 and CA1-2 hypothesized based on cross-sectional data. Our null finding of change in CA1-2 is even more unlikely to result from low power as the observed effect was negative. That said, we may still be missing brain-cognition associations due to limited power. Fifth, low sample size may also lead to unreliable spurious correlations, and such correlations have been pinpointed as potential causes behind disagreeing longitudinal and cross-sectional results from the same sample (Louis et al., 1986). That said, cross-sectional age-associations and longitudinal changes reported here are unlikely to be spurious; the cross-sectional results fit well to a consistent line of previous findings.

One additional possibility is that change in hippocampal subfields is nonlinear. The use of only two measurement occasions in our study precluded the detection of any non-linear effects. Louis and colleagues (1986) have mathematically shown that linearly modeled cross-sectional and longitudinal slopes from the same sample should agree if the age distributions in cross-sectional samples are Gaussian and the change is linear or quadratic nonlinear. Shapiro-Wilk’s test of normality indicated that our age distributions were normal (for both waves *p* > .1). This leaves open the possibility that the disagreement between cross-sectional and longitudinal slopes for hippocampal subfield volumes in our study are due to nonquadratic (e.g., cubic) nonlinearity of actual change. This nonlinearity is supported by both existing longitudinal studies in the field that have more than two measurement occasions. Canada (2021) used an accelerated longitudinal design with a 4- and a 6-year old cohort followed for up to three years. Using this design, they could assess longitudinal change in hippocampal subfield volumes in a sample of children aged 4 to 8. Intriguingly, the authors found a positive change in all subfields investigated (CA2-4/DG, CA1, and SUB) but only between 5 and 6 years of age. In another study using an accelerated longitudinal design, Tamnes and colleagues (2018) assessed a sample of 8-26 year-old participants at 3 measurement occasions two year apart from one another. These authors showed quadratic effects for the CA1 and cubic effects for the SUB. Both of these subfields started off with an initial increase until 13-15 years of age, whereas all other regions showed a linear decrease. In addition to these data, some cross-sectional studies have also found non-linear age-effects in particular in the case of the SUB (Keresztes et al., 2017; Krogsrud et al., 2014; Lee et al., 2014; Tamnes et al., 2014). That said, given that cubic nonlinear effects have been observed for the SUB only, this explanation falls short of accommodating our discrepant findings for the other three subfields investigated.

### 4.2 Developmental change in the subiculum and its implications for memory development

Our results clearly suggest that the SUB undergoes volumetric increase between ages 6 and 10. This needs to be highlighted because of two reasons. First, it is partly in line with previous longitudinal as well as cross-sectional studies, and so far provides the only consistent observation across extant longitudinal studies. Second, in investigations of hippocampal subfield development, the developmental trajectory of the SUB and its association to memory has received little attention as compared to the DG, and the CA regions. As pointed out earlier, the association between change in SUB volume and change in spatial memory performance we observed in this study was weak and should be dealt with care. However, the only change–change association between memory measures and hippocampal subfields in studies with children has been reported for the association of SUB and source memory (Canada et al., 2021). Thus, although we caution against overinterpretation, we do suggest that longitudinal associations between SUB and memory development warrant more attention.

The role of the SUB in memory development has been under-investigated. The limited evidence available for humans have linked SUB structure and function to delayed recall in adolescents (Jeon et al., 2019), to mnemonic specificity in young and older adults (Nash et al., 2021; Stark and Stark, 2017), and to spatial learning in a lifespan sample (Daugherty et al., 2016). In addition, cross-sectional studies of hippocampal subfield in children have reported associations of SUB volume with context (Lee et al., 2014) and source memory (Riggins et al., 2018), whereas another study testing for similar associations in children provided null results (Keresztes et al., 2017).

Even in animal studies, the role of the subicular complex – comprising the presubiculum, the parasubiculum, and the SUB – has received little attention, and its functions remain elusive. Although several studies provided evidence for the involvement of the subicular complex in spatial memory and episodic memory, these studies also have highlighted the large heterogeneity of its cytoarchitectonic properties, cognitive functions, and maturational profile (Aggleton, 2012; Brotons-Mas et al., 2017; Ku et al., 2017; Lavenex and Lavenex, 2013; O’Mara et al., 2009, 2001). For instance, based on cross-sectional histological examination of hippocampi of rhesus monkeys, Lavenex and Lavenex (2013) suggested an early maturing network connecting the SUB with the EC, and a later maturing network connecting the SUB to the CA1. The same data also suggested differential developmental trajectories of presubiculum, parasubiculum, and SUB. In addition, given that the subicular regions and neuronal layers are part of distinct hippocampal networks (Aggleton, 2012), deducing their specific functions is difficult from either lesion or activation studies (O’Mara et al., 2009). Because these regions are – by constraints of technological limits – lumped together in high-resolution MR imaging of human hippocampal subfields, it is highly likely that such heterogeneity will contribute to the available observations (Canada et al., 2021; Tamnes et al., 2018) of non-linear development of the SUB. Moreover, SUB measures in human high-resolution volumetry often include some transitional zone in which both SUB and CA1 cells are present; it is unclear whether this poor specificity may contribute to the observed effects or lack thereof.

### 4.3 Implications for future research on hippocampal contributions to memory development

Based on the results and theoretical consideration presented in this article, we formulate some suggestions for future studies assessing developmental trajectories of hippocampal subfields. First, inclusion of more than two measurement points per individual will allow researchers to detect non-linear change, if they exist. More than three measurement time points per individual will further allow researchers to detect non quadratic non-linearity in change, if these exist. Second, a priori power calculations based on the growing number of related studies may help to decide on necessary sample size to detect both linear and non-linear effects of change (for available tools, see Brandmaier et al., 2015; Zhang and Liu, 2018). Third, and related, longitudinal sample sizes are necessarily constrained by resources available for recruiting and testing participants, MRI hours, as well as manual or semi-automated hippocampal segmentations. Therefore, a viable route to achieve large enough sample sizes may be to combine data in consortia (cf., Walhovd et al., 2018). Complementing these efforts with non-verbal behavioral measures assessing specific hippocampal functions (e.g., variations of the mnemonic similarity task; Stark et al., 2019) and spatial tasks should enhance such efforts across different nations and regions. Fourth, when reporting longitudinal results, reporting cross-sectional results from the same studies may help tease apart cohort and period effects and actual change. Comparing cross-sectional effects in openly available longitudinal datasets to longitudinal change may be a fruitful direction in this regard.

Longitudinal studies are not without methodological challenges (Louis et al., 1986; Raz and Lindenberger, 2011). Beyond their resource needs, they also amplify practice and test-retest effects in performance (Telzer et al., 2018), which have been related to hippocampal subfield volumes in older adults (Bender et al., 2013). Although practice effects are intuitive in the case of behavioral studies, longitudinal MRI measurement are also subject to them. For instance, because motion-artefacts affect both structural (Reuter et al., 2015) and functional (Satterthwaite et al., 2012) MRI measures, and motion is in turn affected by initial exposure to the scanning environment as well as age, estimations of changes in MRI measures are likely to be confounded by both age and number of repeated measurement occasions (Tamnes et al., 2017; see Satterthwaite et al., 2012 for a demonstration of this effect on functional connectivity).

Finally, longitudinal and cross-sectional results may provide information about distinct mechanisms of change. Complex and sudden changes in one’s environment during development, e.g., schooling, may trigger neural changes distinct from ongoing neurodevelopmental change (Brod, Bunge, & Shing, 2017). For instance, intense new types of learning in the first school year may lead to synaptogenesis in DG/CA3 and CA1-2 (Shors, 2004), and perhaps neurogenesis in DG/CA3 (Boldrini et al., 2018; Sorrells et al., 2018), leading to increase in volume, while at the same time, ongoing developmental reorganization of the hippocampal circuit is accompanied by pruning in other hippocampal subfields (Bagri et al., 2003), leading to a decrease in volume. Combining cross-sectional and longitudinal analysis of the same sample may help tease apart such parallel but opposing changes in indirect measures of neural change, such as volumetry.

### 4.4 Limitations

Our results need to be interpreted in light of some limitations: Following extant conventions, we followed a protocol for delineating the EC on six consecutive slices anterior to the hippocampal body (see section 2.2). Using a fix number of slices to extract volumetric measures from at both waves may have biased any estimates of change for EC volume. This makes the observed association between change in EC volume and wave 1 performance on spatial memory challenging. Related to this, due to the low validity of available hippocampal head segmentation methods, our volumetric measures for DG-CA3, CA1-2 and SUB were constrained to the body of HC, therefore we could not capture potential changes in HC head during the observed period. Another limitation of our study is the lack of data collected to allow us to calculate cohort effects, e.g., in which school year participants were. Unfortunately, birth date in this sample is not enough to determine when a child started school. Although cohort effects are unlikely given the small age-range of the sample, schooling at the age of 5-7 has been shown to affect brain function and cognition independent of age (Brod et al., 2017). This result raises the possibility that brain structure may be affected by schooling, even in such a limited age-range. Lastly, additional age-related covariates not assessed in this study may provide a more fine-grained picture of neural changes in the hippocampus. Puberty status may be one such variable of interest as it has been shown to have both a main effect and an interaction effect with sex on hippocampal morphology (Goddings et al., 2014).

## 5. Conclusion

This study highlights striking inconsistencies between cross-sectional age-associations and longitudinal change. Our results specifically question the hypothesis that the DG-CA3, as well as CA1-2 subfields of the hippocampus undergo protracted volumetric increase in middle childhood and that volumetric change in these regions is related to change in memory performance – at least between 6 and 10 years of age. We hope that emphasizing the observed discrepancies, as well as the outlined mechanisms potentially driving them will prove useful to studies of neurocognitive change.

## Acknowledgements

This study was supported by the Jacobs Foundation [grant 2014–1151 to YLS and CH] and conducted at the Center for Lifespan Psychology, Max Planck Institute for Human Development. The work of YLS was funded by a Minerva Research Group from the Max Planck Society, as well as from the European Union [ERC-2018-StG-PIVOTAL-758898], Jacobs Foundation [JRF 2018–2020], and Deutsche Forschungsgemeinschaft (German Research Foundation, [Project-ID 327654276, SFB 1315], “Mechanisms and Disturbances in Memory Consolidation: From Synapses to Systems”). A.K. was supported by the Hungarian National Research, Development and Innovation Office – NKFIH [FK 128648], and a Max Planck Partner Group from the Max Planck Society.

## CRediT author statement

**Attila Keresztes**: Conceptualization, Methodology, Software, Formal Analysis, Investigation, Writing-Original Draft, Writing - review & editing, Visualization. **Laurel Raffington**: Conceptualization, Methodology, Software, Formal Analysis, Investigation, Data Curation, Writing - review & editing, Project Administration. **Andrew R. Bender**: Software, Formal Analysis, Investigation, Writing - review & editing. **Katharina Bögl**: Formal Analysis, Investigation, Writing - review & editing. **Christine Heim**: Conceptualization, Resources, Writing - review & editing, Funding acquisition. **Yee Lee Shing**: Conceptualization, Methodology, Formal Analysis, Writing - review & editing, Supervision, Project Administration, Funding acquisition

