## Supplementary Material for "Longitudinal Developmental Trajectories Do Not Follow Cross-Sectional Age Associations in Hippocampal Subfield and Memory Development"

† Shared last authorship

Corresponding Authors: Attila Keresztes, Research Centre for Natural Sciences, Magyar tudósok körútja 2, 1117, Budapest, Hungary

S1.1 Sex and Age differences between dropouts and non-dropouts per variable

Here, differences were calculated separately for each variable of interest on a subsample having available data for that variable in at least one wave (see Table S1). Forward dropouts included a significantly higher proportion of boys compared to non-dropouts for lure discrimination, $\chi 2(1) = 4.13$, $p =.042$, and hippocampal subfield volumes (for DG-CA3, CA1-2, SUB, $\chi 2(1) = 13.24$, $p < .001$ and for EC, $\chi 2(1) = 11.35$, $p < .001$; note that values differ for EC because data was available for one participant who had no data for the other three hippocampal subfields. No sex differences were observed between backward dropouts and non-dropouts in any variable of interest. Forward dropouts were also significantly older than non-dropouts for associative memory and spatial memory, $t(9.5) = 4.07$, $p = .003$, and hippocampal subfield volumes (for DG-CA3, CA1-2, SUB, $t(40.6)=2.80$, $p = .007$, and for EC, $t(45.3)=2.97$, $p = .004$.). Finally, compared to non-dropouts, backward dropouts were significantly older for associative memory, $t(8.8) = 3.88$, $p = .004$, and significantly younger for lure discrimination, $t(46.1) = 2.14$, $p = .038$.

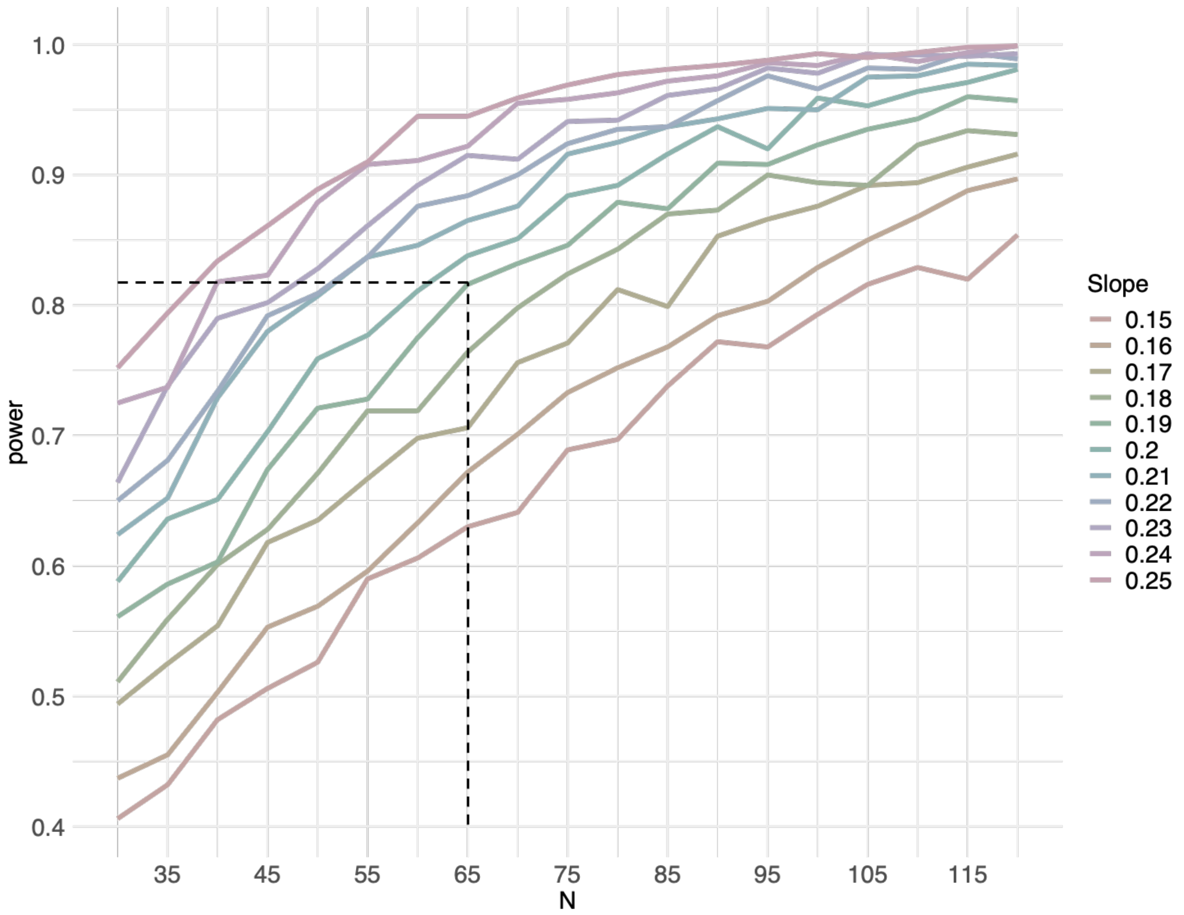

**Figure** S1. Power curves for longitudinal change detection in latent change score (LCS) models for CA1-2 volume. Each line plots power against sample size at a given level of expected longitudinal slope estimate, given an $\alpha = 0.05$. (Note that the smallest cross-sectional slope for CA1-2 across the two waves was 0.34). The dashed vertical line represents the size of our longitudinal sample with complete hippocampal subfield data. The horizontal vertical line points to the last line – representing an expected longitudinal slope of 0.19 –above the conventionally accepted power of 0.8. Power calculations made using RAMpath R package (Zhang et al., 2015; Zhang & Liu, 2018). All parameter values from the actual LCS model were reused as input for the power calculations.

Table S1. Sample size, sex and age descriptives by data collection wave.

| | **Wave 1** | | | | | | **Wave 2** | | | | | | **Wave 1 & Wave 2** | | | | | | | | | |
|---|---|---|---|---|---|---|---|---|---|---|---|---|---|---|---|---|---|---|---|---|---|---|
| | **n** | | **Age (years)** | | | | **n** | | **Age (years)** | | | | **n** | | **Age at Wave 1 (years)** | | | | **Age at Wave 2 (years** | | | |
| | **Total** | **Girls** | **M** | **SD** | **Min.** | **Max.** | **Total** | **Girls** | **M** | **SD** | **Min.** | **Max.** | **Total** | **Girls** | **M** | **SD** | **Min.** | **Max.** | **M** | **SD** | **Min.** | **Max.** |
| **CA1-2Left** | 83 | 40 | 7.32 | 0.41 | 6.08 | 8 | 85 | 33 | 9.24 | 0.44 | 8.34 | 10.12 | 65 | 24 | 7.26 | 0.42 | 6.08 | 8 | 9.27 | 0.42 | 8.38 | 10.12 |
| **CA1-2Right** | 84 | 40 | 7.32 | 0.41 | 6.08 | 8 | 85 | 33 | 9.24 | 0.44 | 8.34 | 10.12 | 65 | 24 | 7.26 | 0.42 | 6.08 | 8 | 9.27 | 0.42 | 8.38 | 10.12 |
| **CA1-2Total** | 83 | 40 | 7.32 | 0.41 | 6.08 | 8 | 85 | 33 | 9.24 | 0.44 | 8.34 | 10.12 | 65 | 24 | 7.26 | 0.42 | 6.08 | 8 | 9.27 | 0.42 | 8.38 | 10.12 |
| **DG-CA3Left** | 83 | 40 | 7.32 | 0.41 | 6.08 | 8 | 85 | 33 | 9.24 | 0.44 | 8.34 | 10.12 | 65 | 24 | 7.26 | 0.42 | 6.08 | 8 | 9.27 | 0.42 | 8.38 | 10.12 |
| **DG-CA3Right** | 84 | 40 | 7.32 | 0.41 | 6.08 | 8 | 85 | 33 | 9.24 | 0.44 | 8.34 | 10.12 | 65 | 24 | 7.26 | 0.42 | 6.08 | 8 | 9.27 | 0.42 | 8.38 | 10.12 |
| **DG-CA3Total** | 83 | 40 | 7.32 | 0.41 | 6.08 | 8 | 85 | 33 | 9.24 | 0.44 | 8.34 | 10.12 | 65 | 24 | 7.26 | 0.42 | 6.08 | 8 | 9.27 | 0.42 | 8.38 | 10.12 |
| **SUBLeft** | 83 | 40 | 7.32 | 0.41 | 6.08 | 8 | 85 | 33 | 9.24 | 0.44 | 8.34 | 10.12 | 65 | 24 | 7.26 | 0.42 | 6.08 | 8 | 9.27 | 0.42 | 8.38 | 10.12 |
| **SUBRight** | 84 | 40 | 7.32 | 0.41 | 6.08 | 8 | 85 | 33 | 9.24 | 0.44 | 8.34 | 10.12 | 65 | 24 | 7.26 | 0.42 | 6.08 | 8 | 9.27 | 0.42 | 8.38 | 10.12 |
| **SUBTotal** | 83 | 40 | 7.32 | 0.41 | 6.08 | 8 | 85 | 33 | 9.24 | 0.44 | 8.34 | 10.12 | 65 | 24 | 7.26 | 0.42 | 6.08 | 8 | 9.27 | 0.42 | 8.38 | 10.12 |
| **ECLeft** | 84 | 40 | 7.32 | 0.41 | 6.08 | 8 | 85 | 33 | 9.24 | 0.44 | 8.34 | 10.12 | 65 | 24 | 7.26 | 0.42 | 6.08 | 8 | 9.27 | 0.42 | 8.38 | 10.12 |
| **ECRight** | 84 | 40 | 7.32 | 0.41 | 6.08 | 8 | 85 | 33 | 9.24 | 0.44 | 8.34 | 10.12 | 65 | 24 | 7.26 | 0.42 | 6.08 | 8 | 9.27 | 0.42 | 8.38 | 10.12 |
| **ECTotal** | 84 | 40 | 7.32 | 0.41 | 6.08 | 8 | 85 | 33 | 9.24 | 0.44 | 8.34 | 10.12 | 65 | 24 | 7.26 | 0.42 | 6.08 | 8 | 9.27 | 0.42 | 8.38 | 10.12 |
| **TotalHC** | 83 | 40 | 7.32 | 0.41 | 6.08 | 8 | 85 | 33 | 9.24 | 0.44 | 8.34 | 10.12 | 65 | 24 | 7.26 | 0.42 | 6.08 | 8 | 9.27 | 0.42 | 8.38 | 10.12 |
| **HairCortisol** | 89 | 46 | 7.25 | 0.44 | 6.07 | 8 | 96 | 45 | 9.27 | 0.44 | 8.34 | 10.16 | 80 | 41 | 7.24 | 0.45 | 6.07 | 8 | 9.28 | 0.43 | 8.34 | 10.16 |
| **LDI** | 73 | 31 | 7.36 | 0.35 | 6.53 | 8 | 96 | 44 | 9.29 | 0.44 | 8.34 | 10.16 | 66 | 25 | 7.37 | 0.37 | 6.53 | 8 | 9.36 | 0.4 | 8.46 | 10.16 |
| **Grid** | 109 | 52 | 7.25 | 0.43 | 6.07 | 8 | 104 | 48 | 9.27 | 0.44 | 8.34 | 10.16 | 104 | 48 | 7.24 | 0.44 | 6.07 | 8 | 9.27 | 0.44 | 8.34 | 10.16 |
| **AMcued** | 100 | 47 | 7.24 | 0.44 | 6.07 | 8 | 103 | 48 | 9.28 | 0.44 | 8.34 | 10.16 | 95 | 43 | 7.23 | 0.45 | 6.07 | 8 | 9.24 | 0.42 | 8.34 | 10.06 |

*Note*. DG: Dentate gyrus, SUB: Subiculum, EC: Entorhinal cortex, HC: Hippocampus, LDI: Lure Discrimination Index, Grid: performance ont he Spatial Memory Task, AMcued: cued recall performance ont he Associative Memory task. At Wave 1, altogether 88 children had gone through a high-resolution hippocampal scan. Of these due to motion, for four children the images were not usable for segmenting hippocampal subfields on either, and for one child on the left hemisphere. At Wave 2, altogether 94 children had gone through a high-resolution hippocampal scan. Of these, due to motion, for nine children the images were not usable for segmenting hippocampal subfields on either hemisphere. Larger dropout due to motion at Wave 2 compared to Wave 1 was probably due to the fact that the high-resolution scan was performed at the end of each session at Wave 2 whereas it was performed in the first half of the scanning session at Wave 1. Hair cortisol data is only available for children who consented hair collection. In addition, in Wave 1, one data point was excluded as an apparent measurement error (>10 SD above mean), and data was below detection limit for another four children. Reasons for missingness for tasks included fatigue and technical errors. LDI has a larger number of missing cases because it was only performed with children attending the MR session, and it was the last task on Day 3.

Table S2. Word pairs used in the associative memory task.

| **Cue word** | **Target word** |
|---|---|
| Topf (Pot) | Esel (Ass) |
| Ofen (Oven) | Heft (Notebook) |
| Mund (Mouth) | Automat (Machine) |
| Tüte (Bag) | Mühlrad (Mill wheel) |
| Bett (Bed) | Trompete (Trumpet) |
| Schublade (Drawer) | Nachbar (Neighbor) |
| Sarg (Coffin) | Bier (Beer) |
| Mantel (Jacket) | Ohr (Ear) |
| Seifenblase (Soap Bubble) | Daumen (Fingers) |
| Burg (Castle) | Maske (Mask) |
| Kiste (Box) | Zauberer (Wizard) |
| Laterne (Lantern) | Roller (Scooter) |
| Boot (Boat) | Schaukel (Swing) |
| Zimmer (Room) | Fischer (Fisher) |
| Schiff (Ship) | Handball (Handball) |
| Helm (Hat) | Bäcker (Baker) |
| Vase (Vase) | Polizist (Policeman) |
| Badewanne (Bathtub) | Giraffe (Giraffe) |
| Mütze (Cap) | Fußboden (Floor) |
| Umhang (Cape) | Wiese (Meadow) |
| Kühlschrank (Fridge) | Schwan (Swan) |
| Flugzeug (Airplane) | Wurm (Worm) |
| Korb (Basket) | Hose (Trousers) |
| Keller (Cellar) | Stein (Stone) |
| Blumentopf (Plant pot) | Sofa (Sofa) |
| Schüssel (Key) | Münze (Coin) |
| Paket (Package) | Eisdiele (Ice cream parlor) |
| Kinderwagen (Stroller) | Wecker (Alarm clock) |
| Auto (Car) | Kreide (Chalk) |
| Käfig (Cage) | Gitarre (Guitar) |
| Mülltonne (Garbage can) | Brücke (Bridge) |
| Schuh (Shoe) | Murmel (Marble) |
| Honigglas (Honey jar) | Brett (Board) |
| Dose (Can) | Pferd (Horse) |

*Note*. For task design and procedure see section 2.4 in the main text. English translations (not used in the experiment) are provided in parantheses.

Table S3. Key parameter estimates in bilateral indicator univariate latent change score models of the hippocampal subfield volumes.

| Variable | Model fit | | | Parameter estimates | | | | | |
|---|---|---|---|---|---|---|---|---|---|
| | | | | $M_{change}$ | | $Var_{change}$ | | $ß_{Age-at-Wave1→Wave1}$ | |
| | $\chi2$ | RMSEA | CFI | PE (SE) | $\Delta\chi2(1)$ | PE (SE) | $\Delta\chi2(1)$ | PE (SE) | $\Delta\chi2(1)$ |
| CA1–2 | 5.07 | 0 | 1 | -0.035 (0.020) | 2.833’ | 0.010 (0.007) | 2.511 | 0.132 (0.064) | 4.258* |
| DG-CA3 | 5.525 | 0 | 1 | 0.015 (0.032) | 0.226 | 0.027 (0.015) | 4.051* | 0.130 (0.109) | 1.399 |
| SUB | 6.78 | 0.034 | 0.994 | 0.12 (0.039) | 8.616** | 0.030 (0.022) | 1.984 | 0.010 (0.12) | 0.008 |
| EC | 6.877 | 0.081 | 0.980 | 0.098 (0.025) | 14.12*** | 0.022 (0.014) | 3.256’ | -0.063 (0.082) | 0.585 |
| Total HC | 2.216 | 0 | 1 | 0.086 (0.085) | 1.024 | 0.188 (0.109) | 3.518’ | 0.273 (0.257) | 1.112 |

*Note*. M: Mean, PE (SE): parameter estimate (standard error), DG: dentate gyrus, SUB: subiculum, EC: entorhinal cortex. Parameters for variances of errors, of change, for covariances between change and Wave 1 values, as well as estimated means of indicator variables are not presented. ’: $p < 0.1$, *:$p < 0.05$, **:$p < 0.01$, ***:$p < 0.001$, uncorrected for multiple comparisons. Confidence intervals calculated from bootstrapped samples provided support for the robustness of all associations significant at $p < .05$. For EC letting variances of left and right indicators differ across waves improved model fit, therefore we lifted these constraints in that model. For model specifications, see Figure 2B.

Table S4. Mean and confidence intervals of bootstrapped parameter estimates that significantly differed from zero in univariate latent change score models

| | Bootstrapped PE | 95% CI |
|---|---|---|
| ***Hippocampal subfields*** | | |
| *Means of change* | | |
| SUB | 0.252 | [0.091,0.414] |
| EC | 0.137 | [0.047,0.227] |
| *Variance of change* | | |
| CA1-2 | 0.171 | [0.119,0.224] |
| DG-CA3 | 0.287 | [0.193,0.381] |
| SUB | 0.556 | [0.327,0.786] |
| EC | 0.155 | [0.103,0.206] |
| HC | 2.567 | [1.673,3.462] |
| *Regression of Wave 1 on Age at Wave 1* | | |
| CA1-2 | 0.345 | [0.028,0.662] |
| *Covariance between change and Wave 1* | | |
| CA1-2 | -0.093 | [-0.158,-0.029] |
| DG-CA3 | -0.199 | [-0.341,-0.058] |
| SUB | -0.428 | [-0.672,-0.185] |
| HC | -1.92 | [-2.95,-0.89] |
| ***Memory measures*** | | |
| *Means of change* | | |
| Cued recall | 0.14 | [0.101,0.179] |
| Spatial memory | 0.082 | [0.049,0.116] |
| LDI | 0.107 | [0.067,0.147] |
| *Variance of change* | | |
| Cued recall | 0.037 | [0.026,0.047] |
| Spatial memory | 0.03 | [0.022,0.037] |
| LDI | 0.025 | [0.017,0.034] |
| *Regression of Wave 1 on Age at Wave 1* | | |
| Cued recall | 0.115 | [0.041,0.19] |
| *Covariance between change and Wave 1* | | |
| Cued recall | -0.014 | [-0.021,-0.007] |
| Spatial memory | -0.013 | [-0.018,-0.008] |
| LDI | -0.017 | [-0.023,-0.01] |

*Note*. See Figure 2a and Table 1 in the main text for model specification, and all parameter estimates of interest, respectively. PE: Parameter estimate, CI: Confidence interval, HC: Hippocampus, LDI: Lure discrimination index.

Table S5. Model fit and parameter estimates for longitudinal parameters of interest in bivariate latent change score models that included bilateral indicator univariate LCS models for hippocampal subfield volumes.

| | **Model fit** | | | **$Cov_{change-change}$** | | **$Cov_{change – wave 1 subfield}$** | | **$Cov_{change – wave 1 memory}$** | |
|---|---|---|---|---|---|---|---|---|---|
| | $\chi2$ | RMSEA | CFI | **PE (SE)** | **$\Delta\chi2(1)$** | **PE (SE)** | **$\Delta\chi2(1)$** | **PE (SE)** | **$\Delta\chi2(1)$** |
| **CA1-2** | | | | | | | | | |
| – LDI | 9.047 | 0 | 1 | -0.002 (0.004) | 0.015 | -0.001 (0.005) | 0.015 | 0.003 (0.004) | 9.752 |
| – Spatial memory | 12.403 | 0.047 | 0.986 | 0.002 (0.003) | 0.388 | 0.001 (0.004) | 0.089 | 0.004 (0.003) | 1.67 |
| – Cued recall | 10.863 | 0.028 | 0.995 | 0.002 (0.004) | 0.104 | 0.002 (0.005) | 0.104 | -0.001 (0.004) | 0.056 |
| **DG-CA3** | | | | | | | | | |
| – LDI | 10.058 | 0.007 | 1 | 0 (0.006) | 0.005 | -0.005 (0.008) | 0.412 | 0.004 (0.006) | 0.421 |
| – Spatial memory | 15.884 | 0.073 | 0.972 | 0.008 (0.006) | 2.067 | 0.001 (0.008) | 0.037 | 0.006 (0.005) | 1.344 |
| – Cued recall | 13.913 | 0.06 | 0.982 | -0.003 (0.007) | 0.237 | -0.002 (0.009) | 0.058 | 0.004 (0.006) | 0.571 |
| **SUB –** | | | | | | | | | |
| – LDI | 11.178 | 0.033 | 0.991 | 0 (0.003) | 0.003 | -0.006 (0.009) | 0.481 | 0.006 (0.007) | 0.85 |
| – Spatial memory | 12.763 | 0.05 | 0.981 | 0.013 (0.007) | 4.135* | -0.005 (0.008) | 0.334 | 0.005 (0.006) | 0.615 |
| – Cued recall | 10.202 | 0.014 | 0.999 | 0.005 (0.008) | 0.327 | -0.002 (0.01) | 0.054 | 0.001 (0.007) | 0.033 |
| **EC** | | | | | | | | | |
| – LDI | 17.437 | 0.083 | 0.952 | -0.004 (0.005) | 0.614 | -0.003 (0.006) | 0.229 | 0.006 (0.005) | 1.318 |
| – Spatial memory | 17.96 | 0.085 | 0.952 | -0.002 (0.004) | 0.37 | 0.002 (0.006) | 0.156 | 0.008 (0.004) | 4.713* |
| – Cued recall | 19.043 | 0.091 | 0.948 | -0.001 (0.005) | 0.082 | 0.007 (0.007) | 1.142 | 0.001 (0.004) | 0.091 |
| **HC** | | | | | | | | | |
| – LDI | 6.67 | 0 | 1 | -0.003 (0.015) | 0.037 | -0.013 (0.019) | 0.46 | 0.015 (0.015) | 0.989 |
| – Spatial memory | 11.475 | 0.037 | 0.99 | 0.022 (0.015) | 2.241 | 0.000 (0.018) | 0.001 | 0.016 (0.014) | 1.347 |
| – Cued recall | 8.629 | 0 | 1 | 0.006 (0.018) | 0.106 | -0.004 (0.021) | 0.042 | 0.003 (0.015) | 0.043 |

*Note*. M: mean, Var: variance, PE (SE): parameter estimate (standard error), DG: dentate gyrus, SUB: subiculum, EC: entorhinal cortex, HC: hippocampus. LDI: Lure discrimination index. $COV_{change-change}$: covariance between change in both variables, $COV_{change–wave 1 subfield}$: covariance between change in memory and hippocampal subfield volume at wave 1, $COV_{change–wave 1 memory}$: covariance between change in hippocampal subfield volume and memory at wave 1, *:$p < 0.05$, uncorrected for multiple comparisons.
